# Cortical Excitability is Affected by Light Exposure – Distinct Effects in Adolescents and Young Adults

**DOI:** 10.1101/2024.08.21.608922

**Authors:** Roya Sharifpour, Fermin Balda, Ilenia Paparella, John Read, Zoé Leysens, Sara Letot, Islay Campbell, Elise Beckers, Fabienne Collette, Christophe Phillips, Mikhail Zubkov, Gilles Vandewalle

**Affiliations:** GIGA-CRC Human Imaging, University of Liège, Liège, Belgium; Alzheimer Centre Limburg, Mental Health and Neuroscience Research Institute, Faculty of Health, Medicine and Life Sciences, Maastricht University, Maastricht, Netherlands; Psychology and Neuroscience of Cognition unit, University of Liège, Liège, Belgium

**Keywords:** Blue-Light, Non-Image Forming, TMS-EEG, Cortical excitability, Adolescents

## Abstract

Light, particularly blue-wavelength light exerts a broad range of non-image forming (NIF) effects including the stimulation of cognition and alertness and the regulation of mood, sleep and circadian rhythms. However, its underlying brain mechanisms are not fully elucidated. Likewise, whether adolescents show a different NIF sensitivity to light compared to adults is not established. Here, we investigated whether cortical excitability, a basic aspect of brain function that depends on sleep-wake regulation, is affected by blue light and whether the effect is similar in young adults and adolescents. We used transcranial magnetic stimulation coupled to high-density electroencephalography (TMS–EEG) in healthy young adults (N=13, 24.2 ± 3.4 y) and in adolescents (N=15, 16.9 ± 1.1 y). Our results showed that, in young adults, blue light affected cortical excitability following an apparent inverted-U relationship, while adolescents’ cortical excitability was not significantly different under blue light compared to orange light. In addition, although light did not affect performance on a visuomotor vigilance task completed during the TMS-EEG recordings, cortical excitability was positively correlated to task performance in both age groups. This study provides valuable insights into the complex interplay between light, cortical excitability, and behavior. Our findings highlight the role of age in NIF effects of light, suggesting that brain responses to light differ during developmental periods.

## 1 Introduction

Besides enabling vision, light exerts a broad range of physiological, hormonal, and neurobehavioral effects not directly related to vision that are often referred to as non-image forming (NIF) effects of light [1,2]. NIF effects are primarily mediated by a subclass of retinal ganglion cells (RGCs) that act as photoreceptors by expressing the photopigment melanopsin [1,2]. These intrinsically photosensitive RGCs (ipRGCs) combine rods and cones outputs to their own response to light and funnel light signal to many (subcortical) parts of the brain involved in the regulation of the circadian system, sleep need, mood and cognition [3–6]. Since melanopsin is maximally sensitive to short wavelength blue light, at around 480 nm, NIF impacts of light on brain functions are mostly driven by the blue-wavelength content of light [7,8]. To reflect the sensitivity of NIF responses, a novel illuminance measurement unit has been defined, i.e. in Melanopic Equivalent Daylight Illuminance (melEDI) which reflects how effective artificial or natural light is at engaging ipRGCs, compared to daylight [9]

A basic aspect of the brain functioning is cortical excitability that can be defined as the responsiveness and response selectivity of cortical neurons [10]. It is therefore fundamental to cognitive brain functions and dictates the impact of an incoming stimulus on cortical activity and on behavior. Cortical excitability depends on prior sleep-wake history and circadian phase [11–13]. It remains relatively stable during the day while well rested and sharply increases if one stays awake[14] such that overnight increase of cortical excitability is correlated to performance decrement [11] and is therefore likely to contribute to the detrimental effect of sleep loss on cognition. The relationship between performance and cortical excitability may follow an inverted U-shape function where an optimal mid-range level of excitability, likely happening during the day when well-rested [15], is associated with optimal performance while lower or higher levels lead to poor cognitive outcomes. Cortical excitability depends on several environmental factors that directly or indirectly affect alertness or sleepiness level, including alcohol and potentially caffeine consumption [16–18]. It is therefore plausible that ambient light could also affect cortical excitability. Findings of a small-scale study, carried out close to 3 decades ago on seven young adults (21-25y), suggested in fact, that there might be an inverted U-shape relationship between light-induced increase in arousal and indirect measure of cortical excitability inferred from an evoked EEG response[19]. Besides this early report, whether exposure to light affects cortical excitability is not established.

Adolescence may be of particular interest when focusing on NIF effects of light. Optimizing the lighting in classrooms has indeed been proposed as a promising simple means to improve alertness and performance at school [20,21]. Research also reported that NIF effects depend on the prior light history, such that prior exposure to high light levels over the preceding hours, days or even weeks, may reduce NIF effects [22,23]. Adolescents are high consumers of LED devices, typically enriched in blue light, and may spend more time outdoors, they may therefore present a reduction in NIF effects of light on brain function. Adolescents, however, are characterized by a late chronotype (i.e. intrinsic time-of-day preference to be physically and/or mentally active), and late chronotypes in adults are typically associated with a higher sensitivity to NIF impact of light [24] that can lead to an increase in NIF effects.

Although it is established that exposure to light affects cognitive brain functions [7], whether spectral composition of light impacts cortical excitability, and whether it happens differently in adolescents versus young adults, is not yet established. Such questions need to be resolved before one can use the flexibility of LEDs to design truly individualized human-centered lighting.

In the present study, we used transcranial magnetic stimulation coupled with high-density electroencephalography (TMS-EEG), as a non-invasive tool to assess cortical excitability in vivo in 13 healthy young adults aged 19-30y while they were exposed to light of different illuminance levels (as computed in Melanopic Equivalent Daylight Illuminance - mel-EDI lux). We further assessed whether cortical excitability was correlated to the performance on a concomitant visuomotor vigilance task. Data were also collected in 15 healthy adolescents aged 15-18y to determine whether sensitivity to the impact of light on cortical excitability was altered in this age group. We hypothesized that in young adults, cortical excitability would be increased by increasing melanopic illuminance and performance on the task would be correlated with cortical excitability. We further anticipated that the impact of light would be different in adolescents with no prior expectation on the directionality of the difference.

## 2 Materials and Methods

The protocol was approved by the Ethics Committee of the Faculty of Medicine of the University of Liège under approval number B7072021000021.

### 2.1 Participants

Between March-2022 and September-2023, thirty-six healthy volunteers participated in the study, 19 adolescents (16.8±1.0 years old, 6 Females) and 17 young adults (24.9±3.9 years old, 9 Females). Adults provided written informed consent, and parental or guardian consent was obtained for underage participants, in accordance with the Declaration of Helsinki. Exclusion criteria were as follows: BMI>28; recent psychiatric history; severe trauma; sleep disorders; addiction; chronic medication; smoking, excessive alcohol (>14 units/week) or caffeine (>3 cups/day) consumption; night shift work during the last year; transmeridian travel during the past 2 months; anxiety [25] or depression [26]; poor-sleep quality [27]; excessive self-reported daytime sleepiness[28]; extreme late/early chronotypes[29]; and pregnancy. We used chronological age to distinguish between adolescents and adults. Assessment of maturation which better reflects developmental status (e.g. Tanner scale) was not included in the study.

Seven participants were excluded from the analyses due to poor data quality, specifically the presence of TMS artefacts that masked the response of interest and were too extensive to be corrected during data processing, and one participant was excluded due to both cortical excitability metrics > 3 standard deviations (SD) compared to the rest of the sample(outlier). Thus, the data presented here includes twenty-eight participants (15 adolescents and 13 young adults). Table1 summarizes the demographic characteristics of the final sample.

### 2.2 Experimental protocol

To avoid excessive sleep restriction while maintaining real-life conditions, participants were asked to keep a regular sleep–wake schedule (±1h), in agreement with their preferred bed and wake-up times, for five days preceding the in-lab experiment. Compliance was verified using sleep diaries and wrist actigraphy (Actiwatch,Cambridge Neurotechnology,UK). Additional methodological information can be found as supplementary method.

On the experiment day, participants arrived at the laboratory between 13:00-14:00. This timeframe was chosen primarily for practical reasons, as this was the period during which adolescent participants were available outside of school hours. For the few sessions conducted during weekends, we maintained the same timeframe to ensure consistency across all participants and minimize potential variability related to time of day. To control for recent light history, participants were maintained in dim-light (<10 lux) for 1:28±0:27, during which the optimal TMS parameters (i.e., location, intensity, coil orientation) providing artifact free TMS-EEG recordings were determined and set. This procedure standardizes participants’ recent light exposure history and minimized any residual effects of prior environmental light exposure (though potentially not eliminating them all). The stimulation target was set to the superior frontal gyrus (SFG) on the dominant hemisphere (left hemisphere as all our participants were right-handed). This brain area was chosen for the following reasons: (1) the SFG is highly sensitive to sleep pressure, including at the neuronal level, just like the entire frontal lobe [30] ; (2) it plays an important role in cognitive performance [31]; and (3) its stimulation does not cause muscle activation, which is a source of EEG signal contamination.

### 2.3 Light set-up and protocol

The light source used in this study was a tunable 35cm×45cm light LED box (EOS,Balder,Huy,Belgium) which allowed us to create three light conditions including orange light served as control light condition, and active lower-intensity and higher-intensity blue light conditions. The illuminance level of each light condition was adjusted manually for each participant such that the orange light and lower-intensity blue light conditions had the same photopic illuminance of 30 lux at the eye level while differing in terms of melEDI illuminance (orange: 24 melEDI lux; blue: 312 melEDI lux). The illuminance for the higher-intensity blue light condition was increased to 60 lux corresponding to 625 melEDI lux. (**Table S1**). **Figure 1** shows the spectrums of the three light conditions.

**Figure 1:**
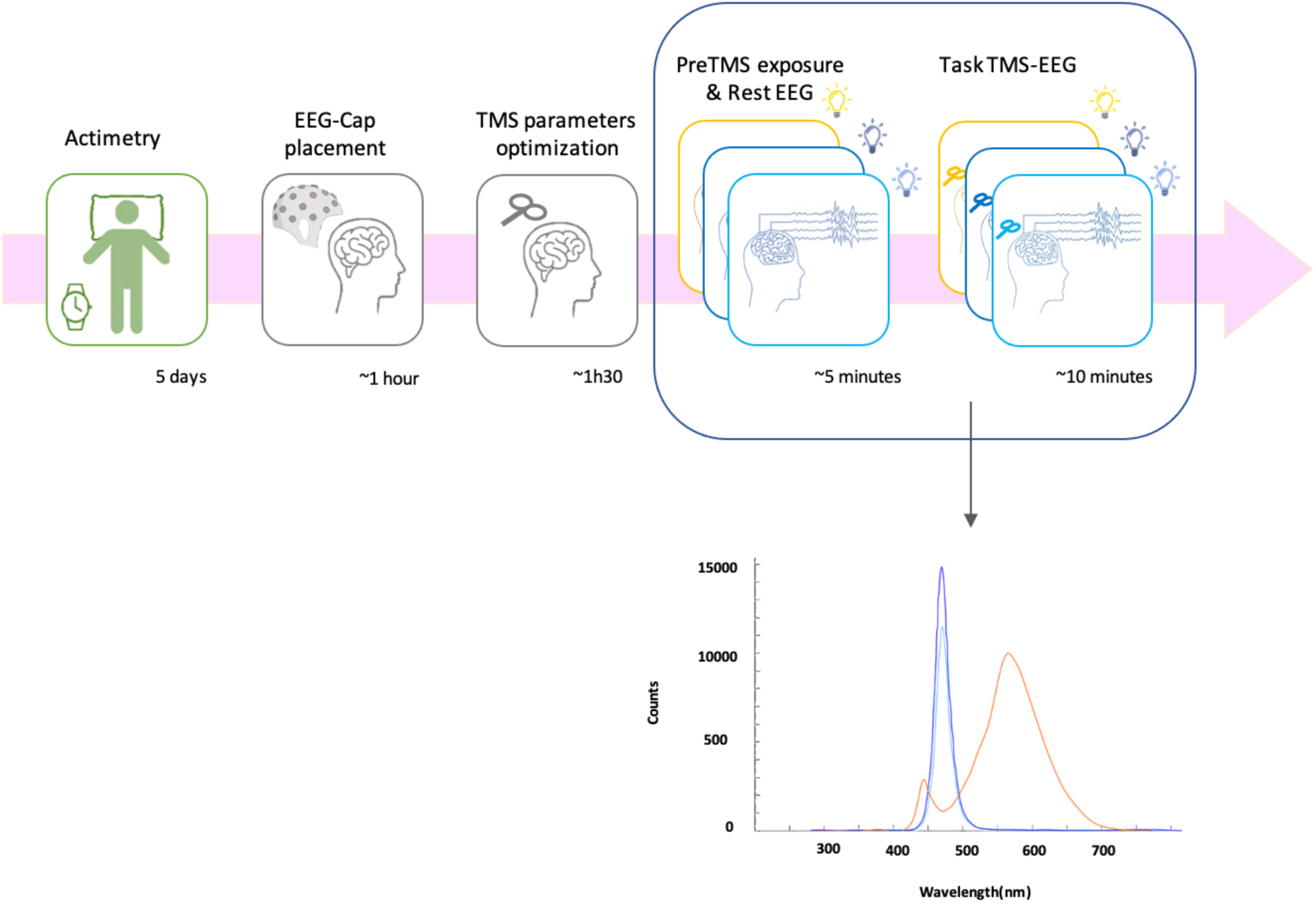
Graphical representation of the protocol and the spectrums of light conditions: Polychromatic orange light at 24 melEDI lux; Monochromatic lower-intensity blue light at 312 melEDI lux, and monochromatic higher-intensity blue light at 625 melEDI lux.

Each light session began with 1-2 mins of light adjustment followed by a two-minute eye-open waking EEG recording. In each light session, TMS-EEG recording pre-exposure time was approximately 5-mins, and the entire TMS-EEG recording time was ∼12 mins. The order of the orange and lower-intensity blue sessions was randomized, with the higher-intensity blue session always being last. The main question of this study was to determine the impact of blue light on cortical excitability compared to the orange light with the same photopic illuminance. Therefore, given the long duration of the entire protocol (around 5 hours) and the possibility of teenage volunteers becoming fatigued and withdrawing early, before the last session of the experiment (happened for one subject), the order of the sessions was decided as explained so that we would anyway have the two sessions needed to answer our main research question. The three sessions were separated by at least a 15-minute washout period in dim light (<10 lux).

### 2.4 TMS-EEG acquisition

A Focal Bipulse 8-shape coil (Eximia; Nexstim,Helsinki,Finland) was used to generate TMS pulses (Suppl.Method). The stimulation target (SFG) was located on individual structural MRI using a neuronavigation system (Navigated Brain Stimulation; Nexstim, Helsinki, Finland).

Each session included around 250 TMS stimulations. The interstimulus interval varied randomly from 1900 to 2200ms. The coil recharging time was set at 900ms after TMS. TMS-evoked responses (TEP) were recorded using a 60-channel TMS-compatible EEG amplifier (Eximia; Nexstim, Helsinki, Finland). The reference and ground electrodes were placed on the forehead. Two extra bipolar electrodes were also used to record the electrooculogram (EOG).

### 2.5 Wake EEG acquisition

Prior to each TMS session, eye-open rest-waking EEG was recorded using the same 60-channel TMS-compatible EEG (+2 EOGs) amplifier. Participants were asked to fixate on a black dot, placed on the light box in front of them, for 2 mins while relaxing and avoiding excessive blinking.

### 2.6 Visuospatial vigilance task

Participants were instructed to perform a compensatory tracking task (CTT) [32] concomitantly to the TMS-EEG recordings. Using a trackball device, the aim was to maintain a constantly moving, randomly positioned cursor on a fixed target in the center of a black background on a computer screen that was set to the lowest brightness (Suppl.Method).

### 2.7 TMS-EEG analysis

TMS-EEG data preprocessing (Suppl.Method) was performed using MNE python package[33] (https://mne.tools/stable/index.html). Independent components of the preprocessed EEG recordings were computed using the fastICA algorithm[34] in order to remove clear TMS-induced artifacts (Suppl.Method).

Cortical excitability was inferred from the amplitude and slope of the first EEG component (0–35 ms) of the TEP measured at the artifact-free electrode closest to the hotspot (i.e. the location with highest TMS-induced electrical field estimated by the neuronavigation system). The latter electrode was always located in the stimulated brain hemisphere. It could vary across participants but remained constant at individual level.

### 2.8 Wake EEG analysis

Power spectral densities were computed using a fast Fourier transform on artifact-free 4s windows, overlapping by 2s, using the Welch’s method (pwelch function in MATLAB 7.5.0) (Suppl.Method). EEG activity was computed over frontal region(F3,F1,Fz,F2,F4,FC5,FC3,FC1,FCz,FC2,FC4,FC6) for theta(4.25–8 Hz) and alpha(8.25-12 Hz) frequency bands over the entire 2-min recording.

### 2.9 Statistical analysis

All statistical analyses were performed with SAS version 9.4 (SAS Institute,Cary,NC,USA). Generalized linear mixed models (GLMM; PROC GLIMMIX SAS procedure) were applied separately in each age group on TEP amplitude, TEP slope, power spectra and task performance as dependent variables according to their distribution (Suppl.Method), with subject (intercept and slope) effect included as a random factor, light conditions (3 illuminance levels) as repeated measures with autoregressive correlation type 1 (ar(1)) and session-order, sex, age, and BMI as covariates. In each age group, the analyses included 4 main dependent variables of interest (amplitude, slope, theta power, alpha power): Benjamini and Hochberg False Discovery Rate (FDR) correction considering 4 independent tests was used to test for significant associations [p<.0125 (for rank 1/4); p<.025 (for rank 2/4); p<.0375 (for rank 3/4); p<.05 (for rank 4/4)]. Direct *post hoc* tests of the main analyses were corrected for multiple comparisons using a Tukey adjustment. In each age groups, additional models included the following potential confound variables separately: chronotype, seasonality, depression and anxiety indices, eye color, season, sleep duration (prior night), daytime sleepiness, sleep quality, insomnia index, subjective sleepiness before and after each session, average habitual caffein and alcohol consumption, time awake before first session, electrode of interest distance from hotspot or alpha/theta power. For simplicity’ sake, GLMMs were first performed on each age group separately and then in another GLMM, age group and illuminance-by-age group interaction terms were included to seek statistical differences between the groups. Semi-partial R^² (R^²) values were computed to estimate the effect sizes of significant fixed effects and statistical trends in all GLMMs [35].

## 3 Results

### 3.1 Cortical excitability increases under moderate blue light exposure and is correlated to performance on the vigilance task in young adults

We extracted cortical excitability metrics as both the amplitude and slope of the earliest EEG response evoked by the TMS pulses (TEP; 0–35ms post-TMS) [13], measured at the electrode closest to the hotspot. Statistical analyses employing both metrics separately as dependent variable revealed a significant main effect of the light condition (p<.02; below FDR multiple comparisons correction), and no main effect of the other covariates (**Figure 2**; **Table 2**). *Post hoc* contrasts showed that compared to the orange light (24 mel-EDI lux), both the amplitude and the slope of the TEP were increased during the lower-intensity (312 mel-EDI lux) blue light exposure (amplitude: t=-3.05, p=.005; slope: t=-2.58, p=.016) but not during the higher-intensity (624 mel-EDI lux) blue light (amplitude: t=-1.11, p=.27; slope: t=-1.00, p= .33). TEP amplitude during the latter 624 mel-EDI lux blue light exposure showed a statistical trend (t=1.94, p=0.06) to be lower compared to 312 mel-EDI lux blue light, while no difference was detected when comparing TEP slope (t=1.58, p=0.13).

**Figure 2.**
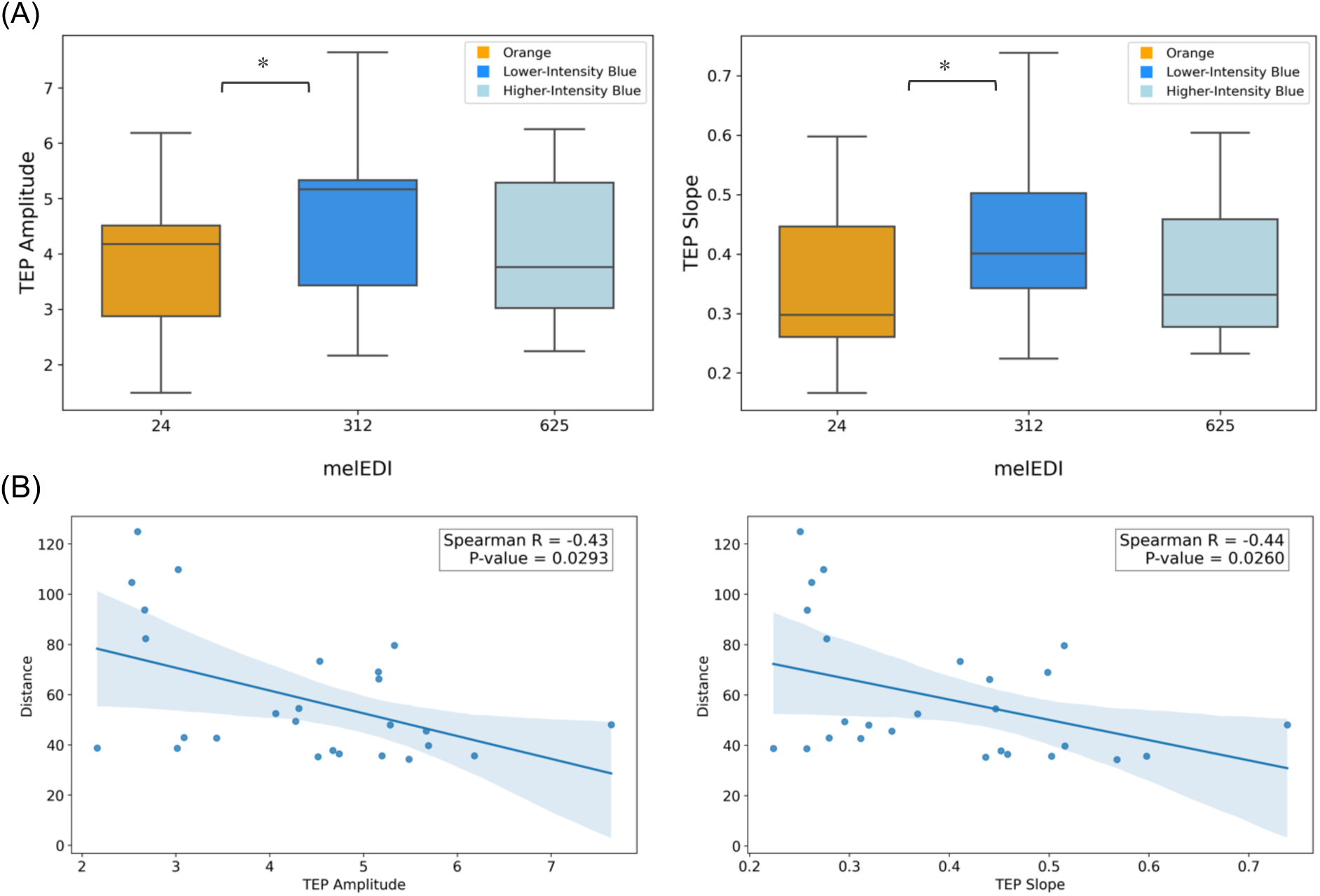
Results in the young adults group. TEP amplitude and slope under different light conditions (A). Regression plots of the association between TEP amplitude and slope with performance on the visuomotor task (B). Spearman R is indicative and do not substitute for GLMM outputs.

**Table 1.** Demographic characteristics of the participants included in the analyses.

|  | Adolescents | Adults | Comparison (t-test) |
| --- | --- | --- | --- |
| Number of Subjects | 15 | 13 |  |
| Age (y) | 16.9±1.1 | 24.2±3.4 | **P<0.001** |
| Sex: Female (Male) | 4 (11) | 5 (8) | P=0.52 |
| Body mass index (kg/m2) | 21.8±2.4 | 22.4±2.3 | P=0.45 |
| Right-handed | 15/15 | 13/13 | P=1.00 |
| Caffeinated drinks (cup/day) | 0.4±0.7 | 1.1±0.9 | **P=0.03** |
| Alcohol (unit/week) | 2.4±4.3 | 2.2±2.0 | P=0.86 |
| Depression (BDI-II<sup>a</sup>) | 7.4±4.7 | 5.1±3.0 | P=0.36 |
| Anxiety Level (BAI<sup>b</sup>) | 6.5±6.0 | 5.9±3.0 | P=0.77 |
| Chronotype (HO<sup>c</sup>) | 42.5±8.3 | 42.6±8.0 | P=0.98 |
| Sleep time (night before the TMS session) | 23:08±0:41 | 23:12±0:46 | P=0.65 |
| Wake time (day of the experiment) | 7:08±0:50 | 7:46±1:09 | P=0.19 |
| Sleep duration (h)<br>(night before the TMS session) | 7.6±1.2 | 8.3±1.2 | P=0.13 |
| Time awake before first session | 9:06±1:25 | 7:30±1:40 | P=0.08 |
| Sleep Quality (PSQI<sup>d</sup>) | 4.3±1.5 | 4.7±1.6 | P=0.83 |
| Habitual daytime Sleepiness<br>(ESS<sup>e</sup>) | 5.9±3.9 | 6.9±2.8 | P=0.45 |
| Insomnia (ISI<sup>f</sup>) | 6.0±3.0 | 6.7±5.1 | P=0.70 |
| Season of experiment * | -0.2±0.6 | -0.2±7 | P=0.90 |
| Seasonality (SPAQ<sup>g</sup>) | 1.1±1.0 | 1.2±0.8 | P=0.83 |
| Eye color: Bright (Dark) ** | 11 (4) | 7 (6) | P=0.30 |
| Electrical Field (Mean±SD) | 115.0±16.7 | 109.3±10.4 | P=0.08 |
| Distance (Mean±SD) *** | 33.0±8.5 | 32.1± 8.9 | P=0.67 |

<sup>a</sup>. Beck's Depression Inventory II
<sup>b</sup>. Beck's Anxiety Inventory
<sup>c</sup>. Horne and Östberg Questionnaire
<sup>d</sup>. Pittsburg Sleep Quality Index
<sup>e</sup>. EPWORTH Sleepiness Scale
<sup>f</sup>. Insomnia Severity Index
<sup>g</sup>. Seasonal Pattern Assessment Questionnaire
\*Cosine (acquisition day of year\*360/365) : 21 December=0°
\*\*Bright: Green and Blue ; Dark: Grey, Brown and Black
\*\*\* Distance (in mm) between the electrode of interest and the stimulation spot (maximum electrical field)
Sleep and wake times assessed based on the actimetry data.
Bold values indicate statistical significance at the p<0.05 level.

**Table 2.** Statistical outcomes of GLMMs with the cortical excitability metrics and theta/alpha spectral power versus the light mel-EDI and covariates.

**Adults**
|  | **Light Condition** | **Session** | **Age** | **Sex** | **BMI** |
| --- | --- | --- | --- | --- | --- |
| **TEP Amplitude** | F(1,23)=8.73<br>**P = 0.007\***<br>**R<sup>2</sup> = 0.27** | F(1,23)=0.46<br>P = 0.50 | F(1,9)=0.50<br>P = 0.50 | F(1,9)=0.05<br>P = 0.83 | F(1,9)=0.05<br>P = 0.82 |
| **TEP Slope** | F(1,23)=6.37<br>**P = 0.019\***<br>**R<sup>2</sup> = 0.22** | F(1,23)=0.00<br>P = 0.98 | F(1,9)=0.80<br>P = 0.39 | F(1,9)=0.22<br>P = 0.65 | F(1,9)=0.01<br>P = 0.93 |
| **Theta Power** | F(1,23)=0.04<br>P = 0.85 | F(1,23)=0.46<br>P = 0.50 | F(1,9)=0.97<br>P = 0.35 | F(1,9)=1.86<br>P = 0.20 | F(1,9)=1.07<br>P = 0.33 |
| **Alpha Power** | F(1,23)=2.96<br>P = 0.10 | F(1,23)=2.82<br>P = 0.11 | F(1,9)=0.52<br>P = 0.49 | F(1,9)=0.16<br>P = 0.70 | F(1,9)=0.55<br>P = 0.48 |

**Adolescents**
|  | **Light Condition** | **Session** | **Age** | **Sex** | **BMI** |
| --- | --- | --- | --- | --- | --- |
| **TEP Amplitude** | F(1,26)=0.09<br>P = 0.76 | F(1,26)=0.02<br>P = 0.90 | F(1,11)=1.12<br>P = 0.31 | F(1,11)=0.01<br>P = 0.93 | F(1,11)=9.36<br>**P = 0.01\***<br>**R<sup>2</sup> = 0.46** |
| **TEP Slope** | F(1,26)=0.01<br>P = 0.92 | F(1,26)=0.05<br>P = 0.82 | F(1,11)=1.44<br>P = 0.25 | F(1,11)=0.11<br>P = 0.75 | F(1,11)=6.23<br>**P = 0.03**<br>**R<sup>2</sup> = 0.36** |
| **Theta Power** | F(1,26)=0.07<br>P = 0.79 | F(1,26)=0.17<br>P = 0.68 | F(1,11)=1.13<br>P = 0.31 | F(1,11)=0.16<br>P = 0.70 | F(1,11)=0.08<br>P = 0.78 |
| **Alpha Power** | F(1,26)=3.01<br>P = 0.094 | F(1,26)=0.67<br>P = 0.42 | F(1,11)=3.12<br>P = 0.10 | F(1,11)=0.04<br>P = 0.85 | F(1,11)=1.28<br>P = 0.28 |

\* Survives per group Benjamini and Hochberg FDR multiple comparisons correction for 4 tests [amplitude, slope, theta power, alpha power; $p < .0125$ (for rank 1/4); $p < .025$ (for rank 2/4); $p < .0375$ (for rank 3/4); $p < .05$ (for rank 4/4)].

Including one of the following potential confounding factors in separate models did not alter the findings, while none of potential confounding factors were significantly associated with cortical excitability (p>0.07): chronotype, seasonality, depression and anxiety indices, eye color, season (in which the sessions were performed), sleep duration (prior night), daytime sleepiness, sleep quality, insomnia index, subjective sleepiness before and after each session, average habitual caffein and alcohol consumption, and time awake before first session; except for anxiety in adults which showed a positive association (p<0.004).

Each TMS-EEG recording was initiated after 5-min of light exposure and lasted about 11 mins (11.5±1.2min). They were preceded by a 2-min recording of the spontaneous EEG activity while quietly awake (1-to-3 mins post-light onset). We computed the power of the EEG over the theta (4.25-8Hz) and alpha (8.25-12Hz) bands as markers of sleepiness/sleep need and alertness, respectively [36,37]. Statistical analyses yielded no significant variation in theta or alpha power across the light conditions (**Table 2**) indicating that baseline brain activity did not differ between the light conditions. Importantly, including theta and alpha power as covariates in our main GLMM did not modify our main statistical outputs (**Table 2**).

In the next step we considered the performance on the visuomotor vigilance task completed during the TMS-EEG recording. All participants performed the task well as indicated by the low number of lapses (1.92±3.1). Statistics indicated that performance was not significantly affected by the light condition (p>0.1) and yet it was significantly correlated to both the amplitude (R=-0.43, p<0.03) and the slope (R=-0.44, p<0.03) of the TEP (**Figure 2**), such that higher cortical excitability was associated with better performance (shorter distance between the moving and fixed dots).

### 3.2 No changes in cortical excitability under different light conditions in adolescents

In adolescents, statistical analyses, using TEP amplitude or slope as dependent variable, did not reveal a significant main effect of light condition (p>0.7) (**Figure 3**; **Table 2**), nor other covariates (except BMI, p=0.03). Including the same potential confounding factors in separate models as in adults did not change the statistical outputs. Likewise, theta and alpha immediately prior to the TMS session did not change across the light conditions (p>0.09; **Table 2**). Again, similarly to adults, including theta or alpha power in the model seeking for an impact of the light condition on the cortical excitability metrics did not modify the statistical outputs of the models (**Table 2**). Examining performance on the visuomotor vigilance task, the statistics revealed similar findings to those found in young adults. Still similarly to adults, performance on the visuomotor was not significantly affected by light condition (p>0.2), but it showed significant correlation with both the amplitude (R=-0.57, p=0.0004) and the slope (R=-0.50, p<0.0028) of the TEPs.

**Figure 3.**
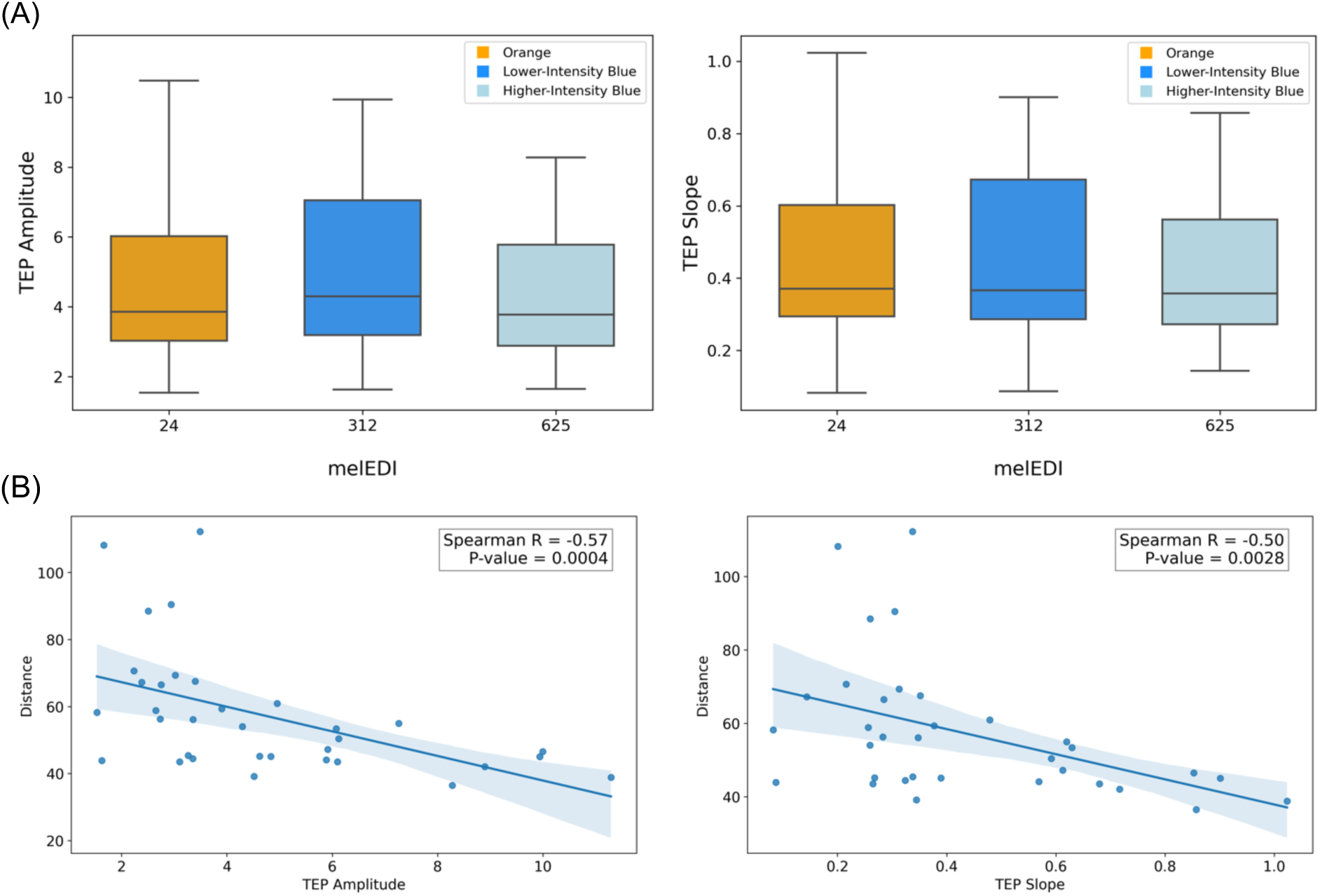
Results in the adolescents group. TEP amplitude and slope under different light conditions (A). Regression plots of the association between TEP amplitude and slope with performance on the visuomotor task (B). Spearman R is indicative and do not substitute for GLMM outputs.

In our final set of analyses, we made a direct group comparison to seek for light condition-by-age group interaction controlling for the same covariates. The GLMM including TEP amplitude as dependent variable yielded a statistical trend for both light condition (F(1,50)=3.81; p=0.056) and light condition-by-age group interaction (F(2,50)=2.77; p=0.072). Conversely, the GLMM including TEP slope as dependent variable revealed no effect of light condition (F(1,50)=2.77; p=0.10) or light condition-by-group interaction (F(2,50)=1.68; p=0.20).

## 4 Discussion

We used TMS–EEG to test whether illuminance affected cortical excitability. Our principal finding indicates that light modulates cortical excitability in humans. We showed that, in healthy young adults, the amplitude and slope of TMS evoked potential increased from lower (∼24 mel-EDI lux) to moderate (∼312 mel-EDI lux) melanopic illuminance, while they may decrease if melanopic illuminance is further increased (∼625 mel-EDI lux). Despite the limited age gap between our two age groups, these effects of light were not detected in adolescents, in which TEPs did not significantly change by light condition. We stress that we did not include maturation assessment while it could better reflect developmental status (e.g. Tanner scale). This constitutes a limitation of our study. Intriguingly, in both age groups, performance on a visuomotor vigilance task was not affected by the light condition and was yet significantly positively related to the amplitude and slope of TEPs. Our findings add to the previously reported multiple NIF effects of light on physiology and behavior and suggest that cortical excitability, which is a basic and fundamental aspect of brain function, may be modulated by environmental light. The data also suggest that the sensitivity of cortical excitability to NIF light effects may differ between adolescence and young adulthood, although further studies are needed to confirm this. Our findings further suggest that optimal cortical excitability changes over different timescales. Further investigations are needed to fully characterize the impact of light at different time of day, with sample of different ages spanning the entire lifetime, and using different light spectra.”

The increase in cortical excitability we detected in young adults between the orange and lower-illuminance blue light can only be attributed to the higher content in blue-wavelength photons of the latter light. Both light conditions were indeed equal in terms of photopic illuminance (in lux) while they differed markedly in NIF illuminance as indexed by the melEDI. This strongly suggests that ipRGC contribute to the increase in cortical excitability we detected. Their projections to many subcortical structures involved in the regulation of alertness (and sleep), including the hypothalamus, may mediate this impact. These subcortical structures would then pass on light stimulating impact to the cortex, either directly, or indirectly through the locus coeruleus in the brainstem [38] or through the pulvinar in the thalamus [39,40] or a combination of multiple subcortical structures. We are, however, in no position to ascertain that only ipRGCs are involved in the effects we detected and stress that other retinal photoreceptors are most likely mediating at least part of the impact of light on cortical excitability, potentially through their inputs to ipRGCs [41].

Our findings in young adults are further compatible with an inverted U-shape relationship between melanopic illuminance and cortical excitability, where increasing illuminance would initially increase cortical excitability before inducing a decrease if illuminance is further increased (though as statistical trends only). This is in line with an early indirect suggestion based on EEG-evoked potential collected in a small sample, relating the impact of light to an effect on the alpha power of the quietly resting EEG [19]. This suggested that alertness change would underlie the inverted U-shape relationship between light illuminance and cortical excitability, such that an optimal range of alertness would be associated with an optimal range of excitability. We did not observe an impact of light illuminance on alpha or theta EEG power and cannot therefore directly relate our findings to changes in alertness or sleepiness. This may be because quiet resting brain activity was recorded after only 1-min of exposure to light which may have been short to induce detectable changes. The impact of light on alertness during the day has in fact not been consistently reported [42–45] and may depend on the study protocol, light quality and the employed technique [46,47].

To guarantee that at least 2 TMS-EEG sessions were completed by all participants, which allowed for testing of our main research question (i.e. does illuminance affect cortical excitability?), the TMS-EEG session with brighter blue light exposure was always performed last. This limitation of the protocol was partly addressed by including session order in our statistical models. We cannot exclude, however, that the trend for a decrease in cortical excitability we observed under high melanopic illuminance is not in part due to a change in time of day. Cortical excitability was indeed reported to decrease in the evening hours, 2-4 hours later than our protocol [11]. In addition, while the pseudo-randomization between the orange and lower illuminance blue light controls for any outlasting effect of an exposure to the next, the last session including the brighter blue light may have been affected by the preceding experiment light conditions. Likewise, although we do not have any indication of this, the brighter blue light exposure might also have triggered more visual discomfort and contributed thereby to the decrease in cortical excitability. Replication of our findings in a larger sample size and a proper session randomization, as well as including other light spectra and illuminance, is therefore warranted to definitely establish an inverted U-shape relationship between melanopic illuminance and cortical excitability.

Contrary to our expectation, in adolescents, cortical excitability was not significantly changed by light illuminance. This could be attributed to either higher or lower sensitivity to light. The theoretical greater transparency of crystalline lens and larger pupil size of teenagers might make them more sensitive to light[48–51] such that a ceiling effect would already be induced by the orange light. However, we did not characterize the crystalline or pupil of our participants. Adolescents are also more prone to be later chronotypes when late chronotype has been suggested to increase the sensitivity to NIF impacts of light [52]. Chronotype is however unlikely to have significantly contributed to our findings as it did not significantly differ between young adults and adolescents in our sample. We note that participants with extreme chronotypes were excluded from the study, such that further research including individuals with extreme chronotypes will be necessary to determine whether the present findings can be generalized to these populations.

Adolescents may also be less sensitive to light such that none of the light conditions we administered affected cortical excitability. Adolescents were likely exposed to more outdoor and/or artificial light over the hours or days preceding the experiment. Despite the standardization of the recent history implemented in the protocol (over the 2h30 preceding the first light exposure), this difference in light history may have reduced their sensitivity to light [22,23]. Given the recommendation for effective light exposure during the day (melanopic lux > 250 melEDI) [53] and the fact that both blue light conditions exceeded this threshold— especially high-intensity blue at 625 melEDI—it seems unlikely that cortical excitability remained unchanged under blue light or reached a ceiling with orange light. We speculate that light may indeed influence cortical excitability under both blue conditions, but due to adolescents’ heightened sensitivity to light, cortical excitability may have fallen at the lower end of the inverted U-shaped function. This could result in no observable differences between the blue light conditions and orange light, as well as between lower-intensity and higher-intensity blue light condition. The inclusion of session recorded in dim-light or in darkness could help determining whether the light exposure we administered induced a significant change in cortical excitability or not.

Still contrary to our expectation, we did not observe a significant impact of melanopic illuminance on performance. Similar to alertness, a significant impact of light on behavior during the day is not always reported and may depend on the exact procedures of the protocols [44,45,54]. The difficulty of the visuomotor task we administered was undoubtedly low such that individual performance might have been (close to) ceiling across the light conditions limiting the potential effect of light. This interpretation is supported by previous neuroimaging investigation of the impact of light which reported significant behavioral impacts using sample sizes larger [55,56] than previous studies reporting no effect using the same task but a smaller sample size [57]. In other words, we speculate that a more difficult task, a longer light exposure and/or a large sample size may lead to the detection of an impact of light in otherwise similar experimental conditions than those used in the present study.

We did, however, find a positive correlation between cortical excitability with vigilance task performance. Performance varied, irrespective of the light condition, in proportion of the variation in cortical excitability. We note that the effect sizes of such associations are stronger than those linking light to changes in cortical excitability. While previous studies reported that the increase in cortical excitability induced by overnight sleep deprivation is associated with poorer performance following a putative inverted U-shape [11], our findings indicate that under well-rested condition in the afternoon, higher cortical excitability is related to better performance.

There may be therefore distinct association between cortical excitability and behavior over different timescales. Sleep deprivation may increase cortical excitability far beyond its optimum level and result in poor performance, while increase in cortical excitability toward an optimum level at a given moment of the normal and well-rested waking day results in improved behavior. These relationships would be equally present in adolescents and young adults. In a smaller time scale, at a given moment of the normal waking day, light exposure, a common environmental factor, can also modulate cortical excitability following an inverted U-shape in young adults and thereby putatively affect behavior, though we provide no evidence in support of this later hypothesis. In adolescents, the impact of light on cortical excitability would be distinct and follows a pattern that remains to be established. This is because of the different sensitivity to light either due to difference in their developing physiology or in light exposure habits. These putative views are summarized in **Figure 4**.

**Figure 4.**
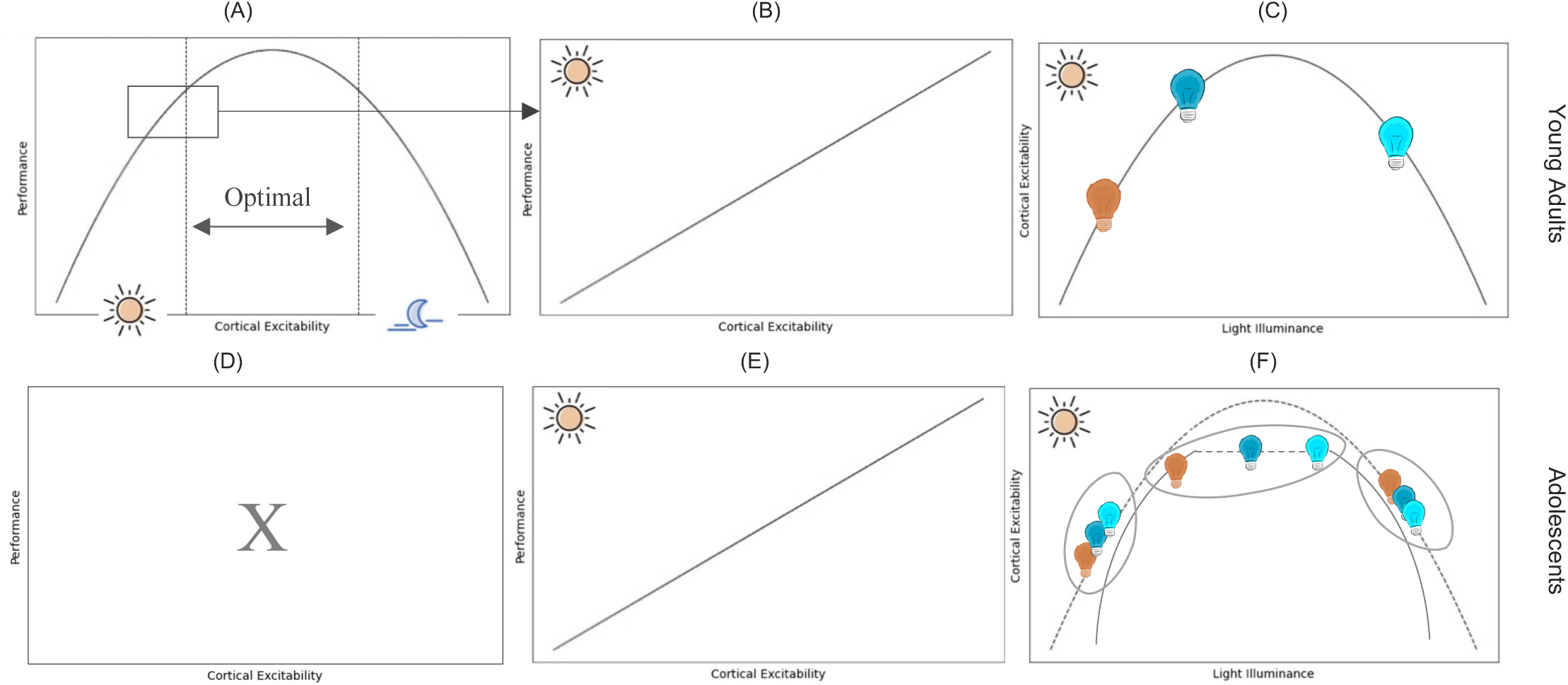
Putative schematic representation of the relationships between cortical excitability, performance and light illuminance in adults (top row) and adolescents (bottom row). (A) In adults, cortical excitability increases with time awake, and correspondence change in performance follows an inverted-U shaped function. Under well-rested condition (current study) higher cortical excitability leads to better performance (B) whereas under extended wakefulness condition, higher cortical excitability has detrimental effects on performance. Our findings suggest an inverted U-shaped relationship between light illuminance and cortical excitability in adults (C). The relationship between cortical excitability and performance is not established in adolescent (D) but our findings confirm a similar relationship to adults under well-rested condition (E). However, our data cannot determine the relationship between cortical excitability and light illuminance in this age group. Some of the possible scenarios have been shown (F).

To conclude, we provide evidence that the quality of environmental light affects brain function down to one of its very basic aspects i.e. cortical excitability. Our findings further suggest that the biological effect of light on cortical excitability is different in adolescents and young adults. Therefore, the development of light interventions in this age group, to alleviate part of their common daytime sleepiness for instance, will need fine-tuning geared toward their specific light sensitivities. The interplay between light, cortical excitability, and behavior leads to complex outcomes and our study lays the foundation for further exploration into the neurobiological effects of light on adolescents and its broader implications for cognitive processes.

## Supporting information

Supplementary materia

## Acknowledgments

The study was conducted at the GIGA-Human Imaging technological platform of ULiège, Belgium. The authors thank Christine Bastin, Christina Schmit, Annick Claes, Christian Degueldre, Catherine Hagelstein, Gregory Hammad, Brigitte Herbillon, Patrick Hawotte, Sophie Laloux, Erik Lambot, Benjamin Lauricella, Pierre Maquet, and Eric Salmon for their help over the different steps of the study.

## Author Contributions

R.S. and G.V. designed the research. R.S, F.B, I.P. J.R., Z.L. and S.L. acquired the data.

All authors contributed to interpretation of the data. R.S. analyzed the data supervised by G.V. R.S. and G.V. wrote the first draft of the paper.

All authors reviewed, edited and approved the final version of the manuscript.

## Data availability statement

The data that support the findings of this study are available from the corresponding author upon request.

