## Supplementary materia for "Cortical Excitability is Affected by Light Exposure – Distinct Effects in Adolescents and Young Adults"

### ONLINE SUPPLEMENTARY MATERIAL

#### Suppl. Method

##### Experimental protocol

Prior to the experimental day, participants went through a structural MRI scan using a 7 Tesla MRI scanner (MAGNETOM Terra, Siemens Healthineers, Erlangen, Germany). A T1-weighted MPRAGE image (TR=2300ms, TE=2.76ms, FA=7°, TI=1050ms, bandwidth=240Hz, FoV=256x256x192 mm$^3$, voxel size=1 mm isotropic spatial resolution) was acquired to use for TMS neuronavigation. Participants were requested to abstain from caffeinated or alcohol drinks and refrain from abnormally intense physical activity for 3 days preceding the study. On the experimental day and to control for recent light history, they stayed in a room equipped with ceiling white LEDs (~180 lux) for 1-hour upon arrival, during which a TMS-compatible EEG-cap was placed. Then they stayed in dim light for almost 1.5h. Three TMS-EEG sessions were performed under different light conditions (**Table S1**), generated by a tunable 35cm×45cm light LED box. The three sessions were separated by at least a 15-minute washout period in dim light (<10 lux) during which participants were allowed to rest while staying on the TMS chair, drink water or have a sugarless snack. Immediately before and after each session, participants reported their sleepiness through the Karolinska Sleepiness Scale (KSS) [1] and rated their motivation, joy, fatigue, stress, anxiety, and effort using the Visual Analogical Scale (VAS).

**Table S1: Detailed characteristics of the light conditions.**

| | *Control Orange* | *Active Low Blue* | *Active High Blue* |
|---|---|---|---|
| ***Illuminance(lux)*** | 30 | 30 | 60 |
| ***Peak Spectral Irradiance (nm)*** | 580 | 470 | 470 |
| ***Melanopic EDI (lux)*** | 23.7 | 312.2 | 625.2 |
| ***Rhodopic EDI (lux)*** | 25.1 | 217.6 | 435.2 |
| ***Cyanopic EDI (lux)*** | 20.3 | 308.7 | 617.3 |
| ***Chlorpic EDI (lux)*** | 28.3 | 106.8 | 213.6 |
| ***Erythropic EDI (lux)*** | 29.1 | 54.1 | 108.2 |
| ***Irradiance (μW/cm²)*** | 9.3 | 44.7 | 89.4 |
| ***Photon Lux (1/cm²/s)*** | 2.63E+13 | 1.06E+14 | 2.12E+14 |
| ***Log Photon Lux ( $log_{10}$ (1/cm²/s))*** | 13.4 | 14.0 | 14.03 |

**TMS-EEG acquisition**

Stimulation target was located on individual structural MRI using a neuronavigation system (Navigated Brain Stimulation; Nexstim, Helsinki, Finland), which allows for exact target localization and reproducible evoked EEG responses (FDA presurgery approval) [2]. The neuronavigation system ensured that hotspot location remained constant across sessions within an individual (±2mm).

The EEG cap was worn by participants throughout the entire protocol, and electrode impedance was kept below 5 kΩ. The signal was bandpass filtered between 0.1 and 500 Hz and sampled at 1450 Hz. The protocol ended with a neuronavigated digitization of each electrode's location.

TMS-induced auditory EEG potentials (AEP) and bone conductance were minimized by playing a continuous loud pink masking noise through earplugs and putting a thin foam layer between the EEG cap and the TMS coil, respectively [3]. Following the last session, 30-40 stimulations were delivered parallel to the scalp in a sham session while the noise was playing at the same level. This session confirmed absence of AEP in all subjects.

**Visuospatial vigilance task**

This task was preferred to psychomotor vigilance task (PVT) during TMS-EEG recordings because it simply requires constant, smooth and limited movement of a single finger and allows for continuous vigilance monitoring [4] . In this task, a lapse was defined as a time during which the cursor remained outside a 200 by 200 pixels box centered on the target for more than 500ms following the last trackball movement. Task performance was calculated as the average distance between the target and moving cursor after excluding lapse periods.

**TMS-EEG analysis**

Continuous EEG recordings were first high-pass and low-pass filtered using an IIR 4th order Butterworth with a cut-off frequency of 1 Hz, and an IIR 15th order Butterworth with a cut-off frequency of 70 Hz, respectively. To minimize the DC offset of the signal, an IIR 4th order Butterworth band-stop filter with a 48-52 Hz range, was applied. Filtered data was first visually inspected to identify and remove bad channels (flat-line or highly noisy/artifactual channels). Individual trials were then split in epochs between 800 to 800 ms post-TMS and bad epochs were rejected during the second run of visual inspection.

The remaining epochs were then re-referred to the average of all good channels. Independent components computed using fastICA, were visually inspected using power spectral density, spatial distribution, and variance distribution over epochs. To minimize data modification, only components representing clear TMS-induced artifacts were set to zero, and the remaining components were used to reconstruct the EEG signals. Artifact free trial epochs were epoched one more time, this time to a shorter period i.e., -300 to 300 ms post TMS. EEG recordings were again successively re-referenced to the average of all good channels and baseline corrected using a window from -101 to -1.5 ms pre-TMS. Epochs were then averaged to have the mean evoked response of each channel in each session.

**Wake EEG analysis**

Waking EEG data were analyzed using MATLAB (2019b, The Mathworks Inc,Natick,MA). Data pre-processing was performed using Statistical Parametric Mapping 12 (SPM12, http://www.fil.ion. ucl.ac.uk/spm). Artifacted channels were rejected after visual inspection. Continuous EEG recordings were re-referenced to the average of all good electrodes and downsampled from 1450 to 500 Hz. Data were then manually and visually scored offline for artifacts (eye blinks, body movements, and slow eye movements).

**Statistical analysis**

Distribution of the dependent variables in the GLMM models were estimated using the allfitdis function in MATLAB (developed by Mike Sheppard, part of the MvCAT package).

**Table S2: Group comparison:** Results of the GLMM analyses examining cortical excitability measures and theta/alpha spectral power in relation to light mel-EDI, age-group and relevant covariates.

**Group Comparison**

| | Light Condition | Age-Group | Light Condition -by- Age-Group | Session | Sex | BMI |
|---|---|---|---|---|---|---|
| **TEP Amplitude** | F(2,50)=3.33<br>P = 0.11 | F(1,24)=1.09<br>P = 0.31 | F(2,50)=2.77<br>P = 0.07 | F(1,50)=0.31<br>P = 0.58 | F(1,24)=0.07<br>P = 0.79 | F(1,24)=6.12<br>**P = 0.02** |
| **TEP Slope** | F(2,50)=1.49<br>P = 0.23 | F(1,24)=0.55<br>P = 0.47 | F(2,50)=1.68<br>P = 0.20 | F(1,50)=0.03<br>P = 0.87 | F(1,24)=0.01<br>P = 0.91 | F(1,24)=4.00<br>P = 0.06 |
| **Theta Power** | F(2,50)=0.09<br>P = 0.91 | F(1,24)=1.62<br>P = 0.22 | F(2,50)=0.96<br>P = 0.39 | F(1,50)=0.62<br>P = 0.44 | F(1,24)=1.71<br>P = 0.20 | F(1,24)=0.98<br>P = 0.33 |
| **Alpha Power** | F(2,50)=2.55<br>P = 0.09 | F(1,24)=0.41<br>P = 0.53 | F(2,50)=0.70<br>P = 0.50 | F(1,50)=0.12<br>P = 0.73 | F(1,24)=0.23<br>P = 0.63 | F(1,24)=1.73<br>P = 0.20 |

**Table S3:** Statistical results from the GLMM analyses assessing performance in relation to light mel-EDI, cortical excitability measures, and relevant covariates in adults and adolescents.

**Adults**

| Light Condition | TEP Amplitude | Light Condition -by- TEP Amplitude | Session | Age | Sex | BMI |
|---|---|---|---|---|---|---|
| F(1,10)=4.24<br>P = 0.07 | F(1,12)=2.69<br>P = 0.13 | F(2,10)=2.01<br>P = 0.18 | F(1,10)=0.38<br>P = 0.55 | F(1,6)=1.81<br>P = 0.23 | F(1,6)=0.27<br>P = 0.62 | F(1,6)=0.66<br>P = 0.44 |

| Light Condition | TEP Slope | Light Condition -by- TEP Amplitude | Session | Age | Sex | BMI |
|---|---|---|---|---|---|---|
| F(1,10)=2.84<br>P = 0.12 | F(1,12)=2.02<br>P = 0.18 | F(2,10)=1.46<br>P = 0.28 | F(1,10)=0.28<br>P = 0.61 | F(1,6)=1.95<br>P = 0.21 | F(1,6)=0.36<br>P = 0.57 | F(1,6)=0.64<br>P = 0.45 |

**Adolescents**

| Light Condition | TEP Amplitude | Light Condition -by- TEP Amplitude | Session | Age | Sex | BMI |
|---|---|---|---|---|---|---|
| F(1,16)=0.26<br>P = 0.62 | F(1,16)=2.29<br>P = 0.15 | F(2,17)=1.55<br>P = 0.24 | F(1,16)=4.34<br>P = 0.054 | F(1,10)=1.68<br>P = 0.24 | F(1,9)=0.60<br>P = 0.47 | F(1,9)=0.31<br>P = 0.60 |

| Light Condition | TEP Slope | Light Condition -by- TEP Amplitude | Session | Age | Sex | BMI |
|---|---|---|---|---|---|---|
| F(1,16)=0.40<br>P = 0.54 | F(1,16)=1.91<br>P = 0.19 | F(2,16)=1.74<br>P = 0.21 | **F(1,16)=4.95**<br>**P = 0.04** | F(1,10)=1.41<br>P = 0.28 | F(1,9)=0.98<br>P = 0.36 | F(1,9)=0.24<br>P = 0.64 |

**Table S4: Group comparison:** Results of the GLMM analyses examining performance in relation to light mel-EDI, cortical excitability ,measures, age-group and relevant covariates.

**Group Comparison**

| Light Condition | TEP Amplitude | Age-Group | Light Condition -by- TEP Amplitude | Light Condition -by- Age-Group | TEP Amplitude -by- Age-Group | Light Condition -by- TEP Amplitude -by- Age-Group | Session | Sex | BMI |
|---|---|---|---|---|---|---|---|---|---|
| F(1,27)=1.85 P=0.18 | F(1,45)=0.01 P=0.91 | F(1,43)=0.03 P=0.85 | F(2,27)=0.64 P=0.54 | F(2,27)=1.40 P=0.26 | F(1,42)=0.21 P=0.65 | F(2,27)=0.85 P=0.44 | F(1,27)=3.67 P=0.07 | F(1,16)=0.41 P=0.53 | F(1,19)=0.05 P=0.82 |
| **Light Condition** | **TEP Slope** | **Age-Group** | **Light Condition -by- TEP Slope** | **Light Condition -by- Age-Group** | **TEP Slope -by- Age-Group** | **Light Condition -by- TEP Slope -by- Age-Group** | **Session** | **Sex** | **BMI** |
| F(1,27)=0.98 P=0.33 | F(1,45)=0.05 P=0.83 | F(1,41)=0.00 P=0.96 | F(2,27)=0.23 P=0.80 | F(2,27)=0.80 P=0.46 | F(1,43)=0.08 P=0.78 | F(2,27)=0.36 P=0.70 | F(1,28)=3.78 P=0.06 | F(1,17)=0.41 P=0.53 | F(1,20)=0.04 P=0.83 |

**Suppl. References**

1. Shahid A, Wilkinson K, Marcu S, Shapiro CM. STOP, THAT and One Hundred Other Sleep Scales. Springer New York; 2012.

2. Rosanova M, Casarotto S, Pigorini A, Canali P, Casali AG, Massimini M. Combining Transcranial Magnetic Stimulation with Electroencephalography to Study Human Cortical Excitability and Effective Connectivity. 2011. p. 435–57.

3. Daskalakis ZJ, Farzan F, Radhu N, Fitzgerald PB. Combined transcranial magnetic stimulation and electroencephalography: Its past, present and future. Brain Res. 2012;1463:93–107.

4. Cardone P, Van Egroo M, Chylinski D, Narbutas J, Gaggioni G, Vandewalle G. Increased cortical excitability but stable effective connectivity index during attentional lapses. Sleep. 2021;44.
